# Volumetric microscopy of CD9 and CD63 reveals distinct subpopulations and novel structures of extracellular vesicles *in situ* in triple negative breast cancer cells

**DOI:** 10.1101/2022.11.08.515679

**Authors:** Elizabeth D. White, Nykia D. Walker, Hannah Yi, Aaron R. Dinner, Norbert F. Scherer, Marsha Rich Rosner

## Abstract

Secreted extracellular vesicles (EVs) are now known to play multifaceted roles in biological processes such as immune responses and cancer. The two primary classes of EVs are defined in terms of their origins: exosomes are derived from the endosomal pathway while microvesicles (ectosomes) bud from the cell membrane. However, it remains unclear whether the contents, sizes, and localizations of subpopulations of EVs can be used to associate them with the two primary classes. Here, we use confocal microscopy and high-resolution volumetric imaging to study intracellular localization of the EV markers CD9 and CD63 prior to EV export from cells. We find significantly different spatial expression of CD9 and CD63. CD9 is primarily localized in microvesicles, while CD63 is detected exclusively in exosomes. We also observe structures in which CD63 forms a shell that encapsulates CD9 and interpret them to be multi-vesicular bodies. The morphology and location within the endoplasmic reticulum of these shell-like structures are consistent with a role in differential sorting and export of exosomes and microvesicles. Our *in situ* imaging allows unambiguous identification and tracking of EVs from their points of origin to cell export, and suggest that CD9 and CD63 can be used as biomarkers to differentiate subpopulations of EVs.

## Introduction

Extracellular vesicles (EVs) is a general term for vesicles that are assembled within a cell and subsequently secreted^1^. After EVs have been secreted from a cell, they are able to travel throughout the body and eventually be taken up by other cells^1–6^. EVs are involved in a variety of systems, including the immune response, cancer, and neurodegenerative diseases^2–5,7,8^. It is commonly accepted that EVs play a critical role in cancer metastasis^8–10^. We have recently shown that EVs secreted by breast cancer tumor cells are able to reprogram macrophages in the environment into pro-metastatic tumor EV-educated macrophages (TEMs)^11^. There is growing interest in using EVs in diagnostic or therapeutic approaches to different diseases^8,12,13^. However, understanding EV function is complicated by the existence of many EV subpopulations. It is theorized that EV subpopulations have distinct roles from one another^14,15^. For instance, a specific EV subpopulation could be involved in cancer metastasis. Identifying EV subpopulations would greatly advance our ability to study EV function.

Two notable types of EVs are exosomes and microvesicles. Both exosomes and microvesicles contain proteins and various forms of RNA^16,17^. Exosomes range from 50-100 nm, and microvesicles range from 50-500 nm^1^. Exosomes are formed in the endosomal pathway^18^. Late-stage endosomes become multivesicular bodies (MVBs) when inward budding occurs to form intralumenal vesicles (ILVs)^18–20^. The MVB can travel to the plasma membrane, fuse the outer membrane to the plasma membrane, and release the ILVs as exosomes ^1,21^. In contrast, microvesicles bud directly from the plasma membrane of the cell^1,22^. Differentiating exosomes and microvesicles is difficult, partially because the size and contents of exosomes and microvesicles are heterogenous and overlapping^1,14,23^.

The most common methods to study EVs are to collect EVs that have been secreted from the cell and isolate subpopulations using ultracentrifugation, density gradients, immunoaffinity assays, or some combination thereof^23,24^. EVs can then be characterized using Western blots, proteomics, or other methods. However, there are significant drawbacks to using *in vitro* methods to study EVs. First, *in vitro* methods separate EVs based on size or density, but they do not provide information on a specific vesicle’s origin. Second, there are some inconsistencies in the terminology in the field, which makes it difficult to precisely define exosome and microvesicle biomarkers^25^. Third, EVs may aggregate during collection or ultracentrifugation, which obfuscates which EV markers are in different populations^14,24,25^.

The transmembrane proteins CD9 and CD63 have been proposed as potential biomarkers to differentiate EV subpopulations. They are both tetraspanins, which are membrane stabilizing proteins^26^. Inhibiting Rab27a, a GTPase that is known to modulate EV secretion^7,27,28^ decreases CD63 export but does not affect CD9 export^27^. This indicates that CD9 and CD63 may be associated with different EV subpopulations and could potentially be used as biomarkers to distinguish the subpopulations^26,28^. However, there is still significant debate about whether CD9 and CD63 are found in exosomes, microvesicles, or both. For instance, some researchers state that CD63 is an exosome-specific marker^12,29,30^, while others state that CD63 is found in both exosomes and microvesicles^14,24^. CD9 is widely reported to be found in both exosomes and microvesicles^10,13–15,31^.

A potentially powerful approach to defining EV subpopulations more precisely is to study EVs intracellularly, prior to export. *In situ* high-resolution imaging can provide detailed information about the spatial locations of small objects, such as vesicles, which is necessary given that EVs are tens to hundreds of nanometers in diameter. Visualizing EV markers intracellularly would provide clearer insight into their origin. In 2021, Mathieu *et al*. visualized and tracked CD9 and CD63 in HeLa cells and saw that CD9 and CD63 were found in two different and one common EV populations^31^. Their work clearly demonstrates that *in situ* imaging is a powerful approach to study EV subpopulations, and that CD9 and CD63 are good candidate biomarkers for distinguishing EV subpopulations.

Volumetric imaging can provide greater insight than traditional 2-dimensional imaging into the morphology and arrangement of vesicles, such as endosomes or MVBs. In other systems, volumetric imaging has been used to reveal endosome sorting and recycling pathways that could not be visualized with 2D microscopy,^32,33^ which demonstrates the advantage of understanding the 3-dimensional structure of vesicles within a cell. Furthermore, *in situ* measurements would allow analyzing the colocalization of potential EV biomarkers in a more biologically relevant context, rather than *in vitro* where the separation between vesicles may be obscured by aggregation. Therefore, high-resolution, volumetric imaging should allow gaining a deeper understanding of where EVs are localized and organized throughout individual cells.

Our work complements and builds off the findings of Mathieu *et al*., 2021. We use high-resolution volumetric imaging to study CD9 and CD63 in triple-negative breast cancer (BM1^34^) cells. Our study reveals that CD9 and CD63 have vastly different intracellular patterns of spatial localization and seldom colocalize, in contrast to previous *in vitro* studies that frequently report CD9 and CD63 colocalizing with one another. The spatial distribution of CD9 and its colocalization with the cell membrane suggest that CD9 is found primarily in microvesicles. CD63, on the other hand, is found only in exosomes. We substantiate this picture with detailed statistical analysis of the colocalization of CD9 and CD63 with one another and with the cell membrane. Finally, we observed 3D structures in which CD63 forms a shell that encapsulates CD9. These shell-like structures are located directly next to the endoplasmic reticulum and appear to be late-stage endosomes or early MVBs involved in EV sorting. These results suggest that CD9 and CD63 can be used to distinguish and characterize distinct subpopulations of EVs. Our work demonstrates that *in situ* imaging of EVs prior to export combined with statistical image analysis is a valuable method that offers unique insights into the subpopulations of EVs.

## Results

### *In vitro* Studies of Extracellular Vesicles Secreted from the Cell Show a High Degree of CD9 and CD63 Colocalization

CD9 and CD63 are often found to be colocalized when EVs are analyzed after being secreted from the cell^13,25^. To verify whether or not this occurs in EVs secreted from triple-negative breast cancer (BM1) cells, we used NanoView technology and quantified marker colocalization (Supplementary Figure 1, see Methods for details). Using this *in vitro* measurement, we observed a high amount of colocalization of CD9 and CD63, which is consistent with other studies that use antibody-based approaches^10,13,16^. These findings for our BM1 cell line are consistent with reported observations for other cell lines.

**Figure 1.**
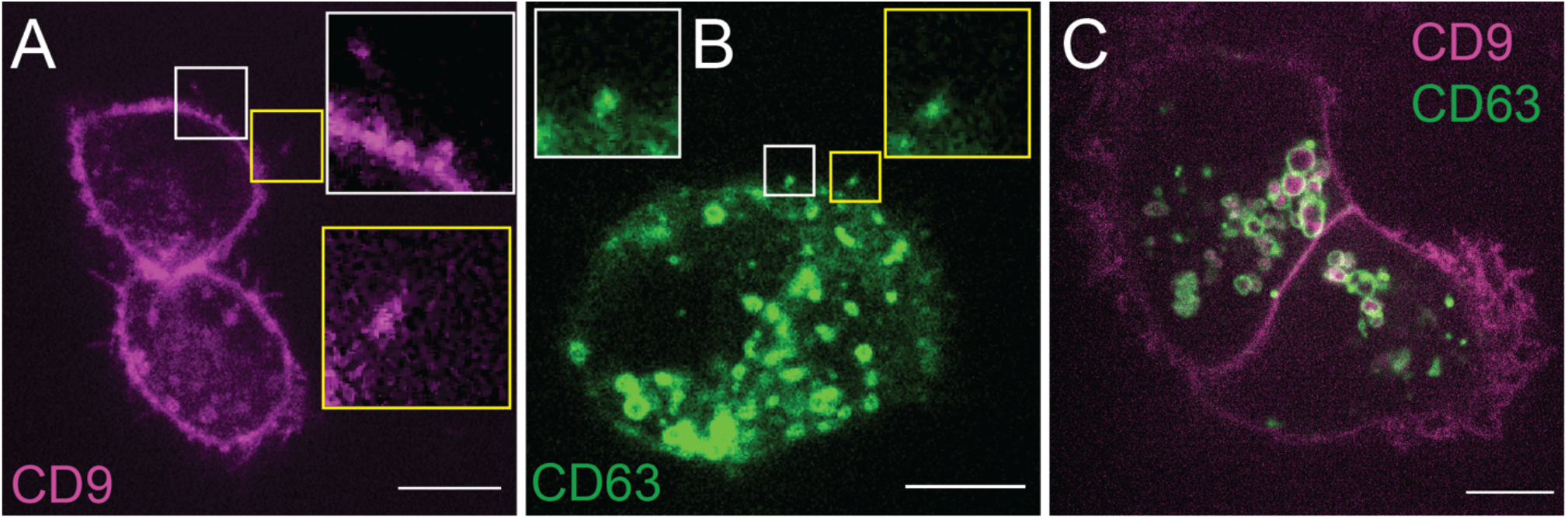
CD9 and CD63 are spatially distinct from one another. BM1 cells were transfected with 0.5 μg CD9-mCherry (left), CD63-eGFP (middle), or both CD9-mCherry and CD63-eGFP (right) and imaged 48 hours post-transfection. Left: CD9 is primarily located at the edge of the cell. Boxes and insets show CD9 EVs being exported from the cell. Middle: CD63 is primarily located in the cytoplasm of the cell. Boxes and insets show CD63 EVs being exported from the cell. Right: CD9 is spatially distinct from CD63. The two proteins are rarely colocalized. Scale bars: 10 μm.

### Intracellular Imaging Shows that CD9 and CD63 Proteins are Spatially Distinct From One Another

We sought to determine whether CD9 and CD63 are truly colocalized intracellularly, or whether the apparent association of these markers might be an artifact of EV isolation. We used volumetric imaging via z-stacks from confocal microscopy to image BM1 cells transfected with CD9-mCherry or CD63-eGFP and observed strikingly different intracellular expression patterns for the two proteins (Supplementary Videos 1 (CD9), and 2 (CD63)). CD9 appeared to be mainly at the cell membrane rather than the interior of the cell (Figure 1A). In contrast, CD63 appeared throughout the cytoplasm in two distinct forms: puncta, or small dots of various sizes, and what appeared to be small, hollow shells (Figure 1B). We observed CD9-containing or CD63-containing vesicles undergoing export from the cell (Figures 1A-B), suggesting these are secreted EVs. CD9 budding was also visualized, but the size of the bud makes it unclear whether these are large microvesicles or small apoptotic bodies (Supplementary Figure 2). To be clear, we did not observe CD63 budding, nor did we observe vesicles with colocalized CD9 and CD63 undergoing export from the cell.

**Figure 2.**
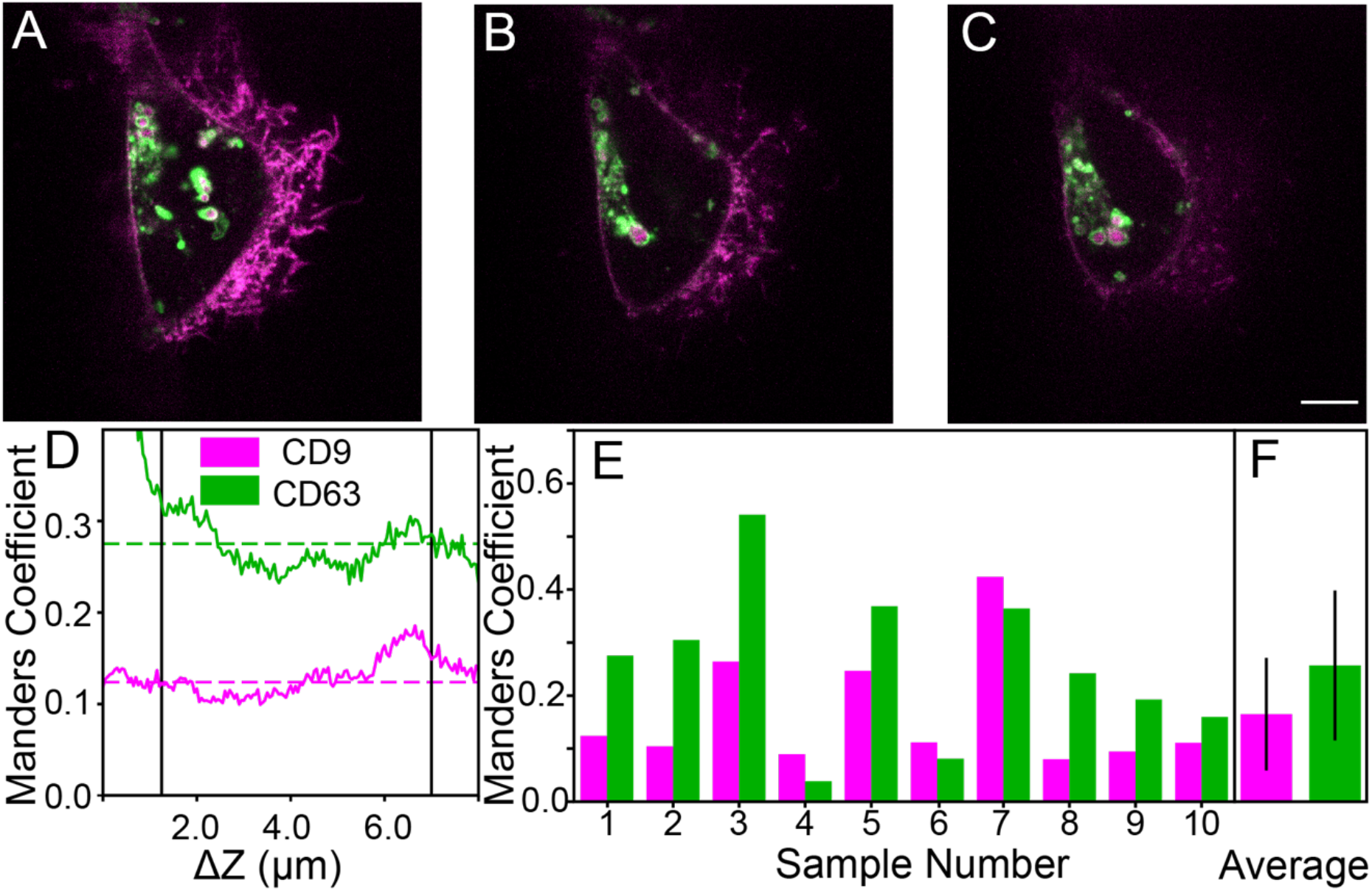
CD9 and CD63 show limited colocalization. Top: BM1 cell transfected with CD9-mCherry and CD63-GFP at (A) 1.25 μm, (B) 3.0 μm, and (C) 5.0 μm above the bottom plane of the cell. Scale bar 10 μm. The cell morphology varies, but the degree of colocalization between the proteins appears to remain limited. (D) The colocalization of the cell was calculated using the Manders coefficient for every medial z-plane (between black lines). The Manders coefficient for CD9 (magenta) and CD63 (green) does not vary significantly over the different medial z-planes of the cell (dotted green line shows average Manders coefficient for CD63, dotted pink line shows average for CD9). (E) The Manders coefficient was calculated for every cell transfected with both CD9 and CD63. (F) The average of the Manders coefficients were calculated for CD9 and CD63 (error bars are standard deviation).

Various concentrations of DNA (0.2 μg-2 μg) were used to transfect cells to determine if the CD9 and CD63 spatial distributions are primarily due to overexpression of the proteins. Regardless of the amount of DNA used, we observed that CD9 appeared localized primarily at the cell membrane and CD63 appeared localized primarily in the cytoplasm (Supplementary Figure 3). In addition, we used immunofluorescence to visualize endogenously expressed CD9 and CD63. We observed a similar trend as described above and conclude that CD9 and CD63 have distinct patterns of localization (Supplementary Figure 4A). To determine the extent of CD9 and CD63 interaction, we transfected cells with both CD9-mCherry and CD63-eGFP (Figure 1c). When observing CD9 and CD63 in the same cell, it became even more clear that there was limited spatial overlap between the two proteins.

**Figure 3.**
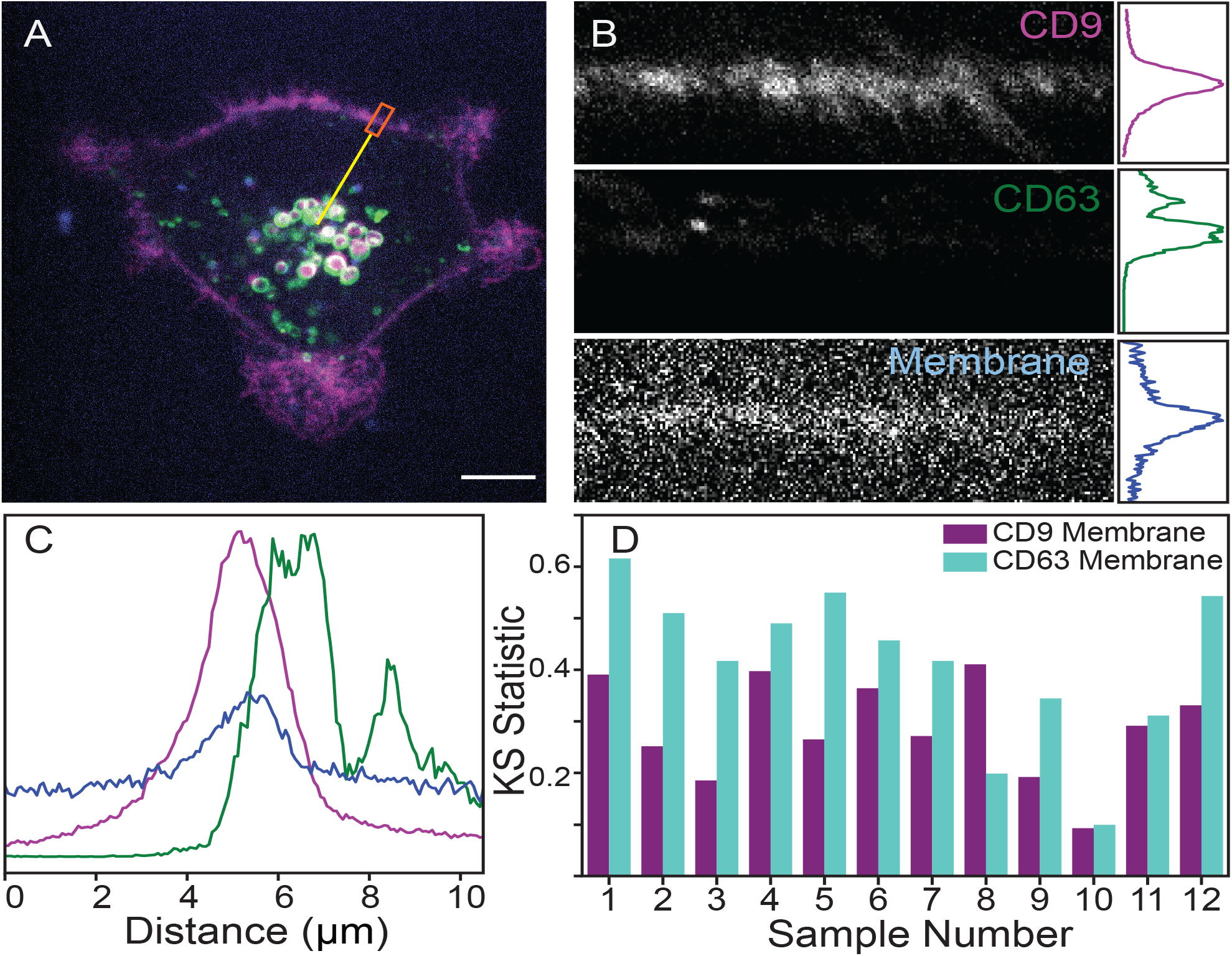
CD9 is more colocalized with the cellular membrane than CD63. (A) BM1 cell transfected with CD9-mCherry (magenta) and CD63-GFP (green), and stained with a membrane dye (blue). Scale bar: 10 μm. Radial line scans were drawn from the center of the cell to the membrane (the yellow line shows an example), and a segment of the line spanning the membrane (orange rectangle) was used to align the radial scans and study the colocalization between proteins and the membrane. (B) Radial line scans across the membrane for CD9, CD63, and the membrane (left). The intensities across all of the radial line scans were averaged (right). (C) The average intensities of the three channels from (B) superimposed. (D) The Kolmogorov-Smirnov test was calculated for regions of the membrane for 10 cells. The KS statistic for CD63 and the membrane was generally greater than the KS statistic for CD9 and the membrane, indicating that the CD9 and membrane have more similar distributions than CD63 and the membrane. Note: Larger KS statistic indicates less similar distributions.

**Figure 4.**
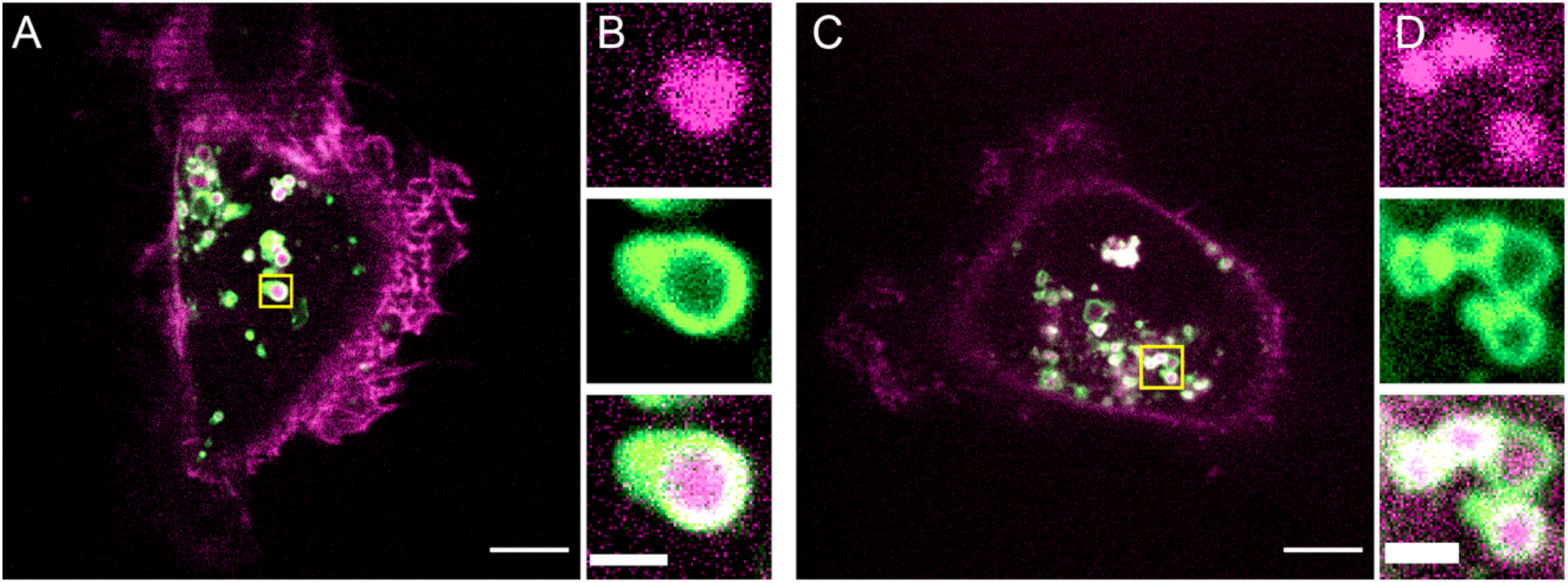
CD63 forms shell-like structures encasing CD9. (A), (C) BM1 cell transfected with CD9-mCherry and CD63-GFP. Scale bar: 10 μm. (B), (D) Expanded views of yellow boxes in (A), (C). Top: CD9, middle: CD63, bottom: composite. Scale bars: 2 μm.

We sought to quantify the colocalization of CD9 and CD63. To do so, we used the Manders coefficient, a statistical test to determine the colocalization of two samples^35,36^. However, cells are 3D systems, and two markers may appear to be colocalized in 2D imaging if they are in the same xy location but different z locations. Therefore, we calculated the Manders coefficient of CD9 and CD63 for each z-plane in a cell (see Methods for details) (Figure 2). We limited this analysis to the medial z-planes of each cell (Figure 2D, area within black bars) to avoid artifacts associated with proximity to the top and bottom of the cell. Using this method, we calculated the percentage of the total CD9 that was colocalized with CD63, and the percentage of the total CD63 that was colocalized with CD9 for all the cells co-transfected with both proteins (Figures 2E-F). Although the results varied from cell to cell, our *in situ* imaging established very limited colocalization between CD9 and CD63. Only 16% of the total CD9 population is colocalized with CD63, and only 25% of the total CD63 population is colocalized with CD9.

To establish that the colocalization patterns were not due to overexpressing CD9 and CD63, we repeated the same analysis in cells with endogenous CD9 and CD63 immunostained with anti-CD9 and anti-CD63 antibodies (Supplementary Figure 4B). Strikingly, we observed even less colocalization than we did with the transfected cells. Therefore, the low level of intracellular colocalization between CD9 and CD63 was consistent regardless of whether the proteins were endogenously expressed or if the cells had been transfected with fluorescently labeled CD9 and CD63.

### CD9 is Significantly Colocalized with the Cell Membrane while CD63 Exhibits Limited Colocalization

CD9 appeared to be primarily at the cell membrane which suggests that CD9 may be used as a marker for microvesicles. Visualization of the membrane and its potential colocalization with CD9 or CD63 would help determine if CD9 is truly colocalized with the cell membrane and if CD9 can be used as a marker for microvesicles. We thus co-transfected cells with CD9-mCherry and CD63-eGFP and stained the membrane with MemBrite 405/430 dye (Figure 3A). Qualitatively, we observed that CD9 was significantly colocalized with the membrane while CD63 was not. We further verified this by quantifying the colocalization of CD9 and CD63 with the cell membrane.

We wanted to measure the colocalization of CD9 and CD63 with the membrane over the entire contour of the membrane. To do so, we drew radial scans from the cell centroid to the cell membrane to collect scans perpendicular to the cell membrane at every location along the membrane contour (see yellow to white radial lines in Figures 3A-B, left). We centered the membrane in each line scan, so we were able to average the line scans from a given cell together to find the spatial distributions of CD9, CD63, and the cell membrane label across the membrane contour (Figures 3B-C). We used the Kolmogorov-Smirnov (KS) test, which is sensitive to both the location (i.e., mean value) and shape of different distributions,^37^ to quantify the colocalization of CD9 and CD63 with the cell membrane for each cell. It is important to note that the KS test measures the difference between two distributions, not the similarity. When two distributions are very similar, the value of the KS coefficient is small; when two samples are very different, the KS coefficient is near 1. For every cell we studied, the KS coefficient of CD9 with the membrane was smaller than the KS coefficient of CD63 with the membrane. Since there was significant variability between cells, averaging of the KS statistics of the cell populations is less useful than applying this test to individual cells. This statistical result demonstrates that CD9 significantly colocalized with the cell membrane, in contrast to CD63, which did not.

### CD63 Forms a Shell That Encapsulates CD9

We found a distinctive structure in which CD63 formed a shell that encapsulates CD9 in the vast majority of cells (Figure 4). Although the CD9 and CD63 signals appeared to be colocalized in 2D measurements, our 3D approach revealed that CD9 and CD63 were not colocalized but in close proximity. CD9 and CD63 occasionally appeared colocalized when the z-plane was at the top or bottom of the shell as a result of the finite (∼800 nm) depth of the confocal slices. To verify that these shell-like structures are not an artifact of overexpression, we confirmed that the same structures are observed in endogenously-expressed protein immunostaining data (Supplementary Figure 5).

**Figure 5.**
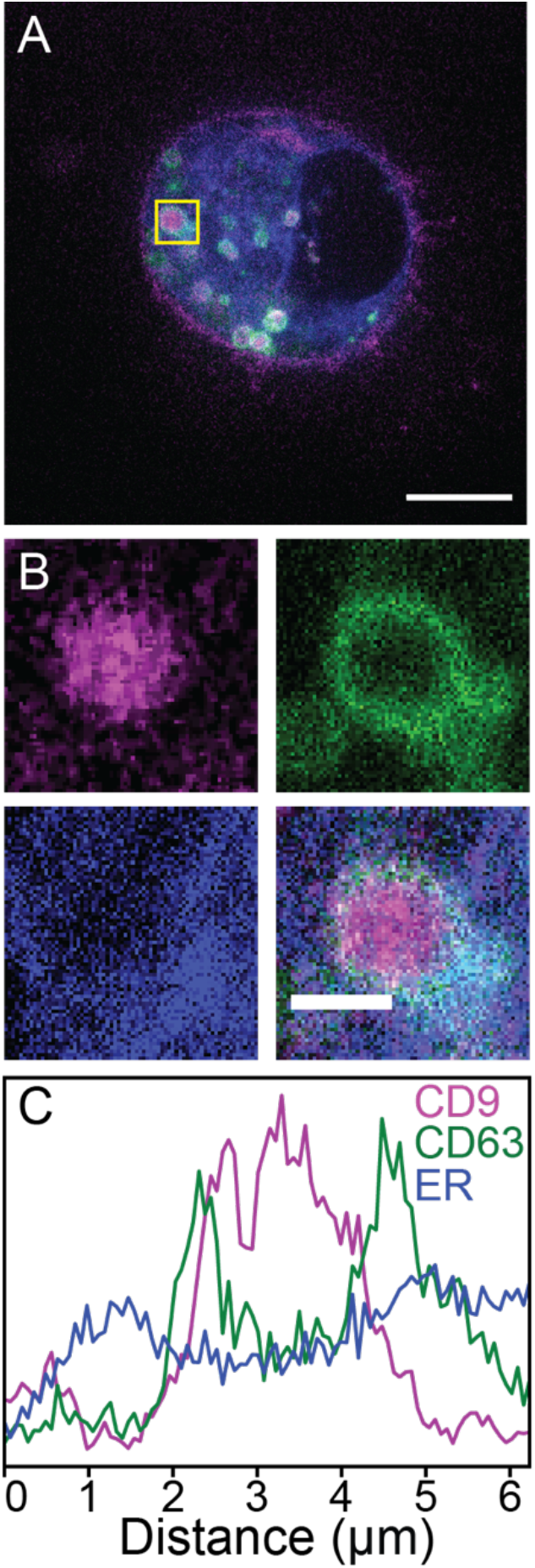
Shell-like structures are found in regions of sparse ER signal, and CD63 is in contact with the ER while CD9 is not. (A) Cells were transfected with CD9-mCherry (magenta), CD63-GFP (green), and BFP-KDEL to label the ER (blue). Scale bar: 10 μm. (B) Zoom in on yellow box in (A), highlighting shell-like structures. Top left: CD9, top right: CD63, bottom left: ER, bottom right: composite. Scale bar: 2 μm. (C) Average intensities for 20 line scans through the center of the shell-like structure for CD9 (pink), CD63 (green), and ER (blue).

### CD63 is Contiguous with the Endoplasmic Reticulum While CD9 is Not

A possible explanation of the shell-like structures we observe is that they are multivesicular bodies (MVBs), which are late-stage endosomes that contain smaller vesicles. MVBs are formed from late endosomes near the endoplasmic reticulum (ER). When there is inward budding from the endosomal membrane, intralumenal vesicles (ILVs) are formed, thus making the late-stage endosome an MVB^18^. MVBs are a sorting mechanism in the cell; ILVs can be marked for degradation, transported to different areas of the cell, or even exported from the cell as exosomes^1,16–18,29^. We considered the possibility that the shell-like structures we observe are MVBs where CD63 is in the MVB-limiting membrane and CD9 is in ILVs within. CD63-associated EVs are known to be found in MVBs^18,38^, but there have not been similar reports about CD9 localizing in MVBs.

Since late-stage endosomes and MVBs are known to frequently have contact sites with the ER^39^, we sought to determine the spatial relationship with CD9, CD63, and the ER. We co-transfected cells with CD9-mCherry, CD63-eGFP, and BFP-KDEL, an ER biomarker^40^ (Figure 5A). We observed holes or voids in the BFP-KDEL signal, and the shell-like structures were frequently found near these holes (Figure 5B). The proximity of the shell-like structures to the ER suggests that they are part of the endosomal pathway. To study this further, we took line scans across the shell-like structures and ER directly adjacent to the CD63-coated shell and compared the distributions of CD9, CD63, and the ER marker (see Methods for details). The line scans shown in Figure 5C suggest that CD9 forms a central core that is surrounded by CD63 that in turn is surrounded by the ER.

## Discussion

CD9 and CD63 were previously reported to share a large degree of colocalization in *in vitro* studies^20,28,29^. For instance, Kowal *et al*. used antibody-coated beads to isolate EVs with different surface markers and used ultracentrifugation to identify different fractions and reported that CD9 and CD63 are found in the same fractions, albeit with varying expression levels^15^. In our own *in vitro* measurements, we observed 33-57% colocalization of CD9 and CD63. However, *in vitro* methods are not sufficiently precise to determine whether CD9 and CD63 can truly be used as biomarkers. There is a strong chance that distinctions between individual EVs, and thus EV subpopulations, is obscured due to EVs aggregating when they are collected or during centrifugation^24,25^. Additionally, *in vitro* methods cannot accurately determine an EV’s origin.

To address these issues, Mathieu *et al*. recently imaged CD9 and CD63 *in situ* in HeLa cells. In that work, they observed that CD9 was localized primarily at the cell membrane and was occasionally found intracellularly^31^. In contrast, CD63 was found primarily intracellularly. They also observed that CD9 and CD63 transiently traffic through MVBs and the cell membrane^31^. They concluded that while CD9 and CD63 can be found in both exosomes and microvesicles, CD9 is primarily in microvesicles while CD63 is primarily in exosomes^31^. Our work complements and builds on Mathieu *et al*. Our studies are in triple negative breast cancer cells, so they provide information about the behavior of these markers in a second cell line. More importantly, our volumetric imaging allows us to dissect the nature of the *in situ* colocalization of CD9 and CD63.

Consistent with the findings from Mathieu *et al*., we observed that CD9 and CD63 had distinct spatial expression patterns. CD9 was primarily found at the edge of the cell, while CD63 was found primarily intracellularly. Furthermore, CD9 was highly associated with the cell membrane, while CD63 was not. This indicates that CD63 is primarily in exosomes, while CD9 is frequently found in microvesicles (Figure 6) in triple negative breast cancer cells, as in HeLa cells.

**Figure 6.**
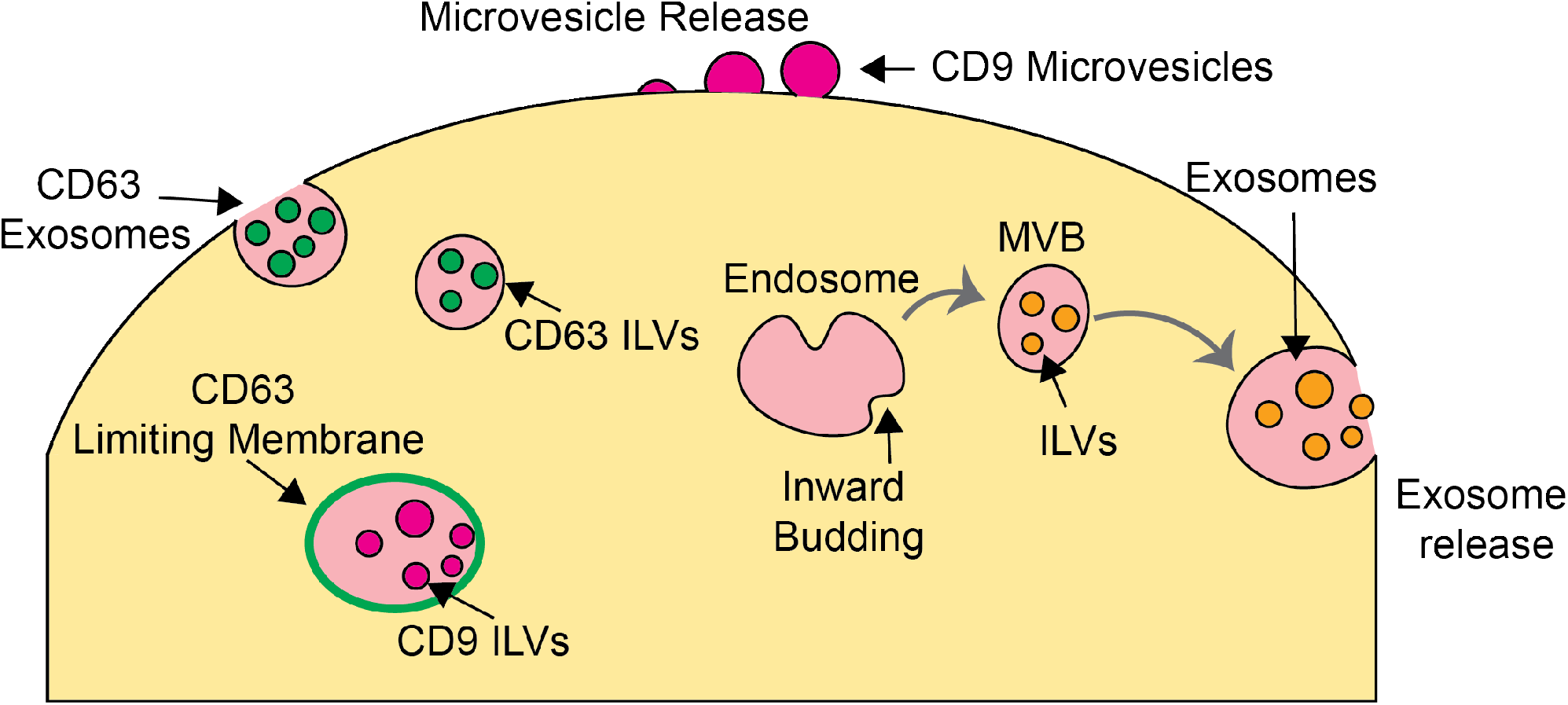
CD9 and CD63 are found in unique populations and structures. CD9 was enriched in the cell membrane and budded from the membrane, likely as microvesicles. Early endosomes stem from the Endoplasmic reticulum. When there is inward budding, interluminal vesicles (ILVs) are formed, and the overall structure is considered a late endosome or multivesicular body (MVB). When the MVB fuses with the membrane and releases the vesicles within, the released vesicles are considered exosomes. We observed puncta with CD63 in the cytoplasm, which were likely CD63-containing ILVs that are waiting to be exported as exosomes. The shell-like structures we observed were likely MVBs in which CD63 was enriched in the limiting membrane, and CD9-containing ILVs were inside the MVB. Note: CD63 and CD9 are found on the membrane of vesicles, not the interior, the coloring was chosen for clarity.

Our volumetric imaging revealed previously unobserved shell-like structures in which CD63 encapsulates CD9. In these shell-like structures, CD63 is contiguous with the ER, while CD9 is not. Based on their structure and association with the ER, we believe these shell-like structures are MVBs in which CD63 is in the outer membrane and CD9 is in ILVs within the MVB (Figure 6). Notably, Mathieu *et al*. saw that CD9 and CD63 transiently colocalize in MVBs. However, they were using 2D imaging and thus did not have the ability to resolve the shell-like structures. In contrast, we can see that while CD9 and CD63 are in the same MVB, they are in different components of the structure. This indicates that they are likely trafficked differently from these shell-like structures.

Quantifying our imaging with the Manders coefficient, we conclude that there is 16-25% overlap in the CD9 and CD63 populations. This establishes statistically that CD9 and CD63 have limited colocalization. Moreover, it is important to note that the shell-like structures constitute a significant portion of the apparent colocalization between CD9 and CD63. The z-plane resolution is such that CD9 and CD63 at the tops and bottoms of the shell-like structures appear to be colocalized, when in reality they are not. Therefore, the Manders coefficient overestimates the true amount of colocalization. This makes the difference in the colocalization between the CD9 and CD63 populations that we observe *in situ* compared to reports of *in vitro* measurements^14,15,29^ even more striking. Likewise, statistically comparing the spatial relationship between CD9 and CD63 with the cell membrane increased our confidence that CD9 is predominantly in microvesicles, while CD63 is not.

Observing EVs *in situ* prior to export provides key insights into distinct populations of EVs. This can be used to advance our understanding of EV origin and function. In the future, tracking EV markers in 3D over time^41^, both intra- and extra-cellularly, could provide a deeper understanding of how EVs are formed and exported from the cell. Sufficiently precise tracking of EV subpopulations could reveal differences in EV type and cargo that impact their ability to reprogram target cells, such as EVs that promote metastasis in cancer cells.

## Methods

### Cell Culture

MDA-MB-231 1833 (referred to as BM1) cells were obtained from Andy Minn^34^, and all work was done within 18 passages of initially establishing the cells. Cells were cultured in DMEM with 10% fetal bovine serum, 50 U/ml penicillin and 50 U/ml streptomycin.

### Preparation for NanoView Assay

BM1 cells were grown to 80% confluence in DMEM media with 10% fetal bovine serum, 50 U/ml penicillin and 50 U/ml streptomycin. Cells were then washed with PBS and incubated in serum-free media for 24 hours. The cell media was centrifuged at 300 g for 5 minutes to harvest the supernatant, containing EVs. The EV concentration was measured using NanoSight technology, and samples were diluted to a final concentration of 2.5e8/mL.

### ExoView Tetraspanin Assay

Samples were run according to the manufacturer’s instructions for the Human Tetraspanin Kit (EV-TETRA-C, NanoView Biosciences Inc.). In brief, samples were diluted 3 times in Solution A and 35 µLs were incubated on the ExoView chips overnight with capture antibodies against human CD81, CD63, and CD9. After sample incubation, chips were washed and stained for 1 hour with ExoView tetraspanin labeling antibodies that consist of (anti-CD81 CF555, anti-CD63 CF647, and anti-CD9 CF488). After staining the chips were washed, dried and imaged with the R100 reader. NanoViewer Analysis software 2.9.1 was used to calculate the particle count and colocalization for each capture spot.

### Transfection

Cells were plated at a ∼2.75e5 concentration on 6 well plates 24 hours before transfection. Cells were transfected using a Lipofectamine 3000 kit following manufacturer’s directions. 0.5 μg CD9-mCherry (Addgene plasmid #5013; http://n2t.net/addgene:55013; RRID:Addgene_55013) and CD63-eGFP (Addgene plasmid #62964; http://n2t.net/addgene:63964; RRID:Addgene_62964) was used unless noted otherwise. 2 μg BFP-KDEL (Addgene plasmid #49150; http://n2t.net/addgene:49150; RRID:Addgene 49150) DNA was used to label the ER. To dye the membrane, cells were incubated with the MemBrite pre-staining solution for 5 minutes at 37° C, then incubated with MemBrite Fix 405/430 (30092-T) for 5 minutes at 37° C. Cells were incubated with transfection reagents for 24 hours post transfection. They were plated on glass bottom Matek 35 mm cell sample chambers at ∼50% confluence. Cells were incubated on these plates for ∼24 hours. Imaging occurred ∼48 hours post-transfection.

### Immunofluorescence

Cells were plated on Matek 35 mm cell sample chambers at ∼50% confluence 24 hours before imaging. Primary antibodies were incubated overnight with fixed and permeabilized cells at 4° C. The next morning, the cells were washed, and the secondary antibodies were added and incubated for one hour at room temperature. Antibodies used: CD9 primary: Thermo Fisher Scientific MA5-33125. CD9 secondary: Thermo Fisher Scientific A32733. CD63 conjugated primary and secondary: Thermo Fisher Scientific MA119602.

### Imaging

Cells were plated on Matek 35 mm cell sample chambers at ∼50% confluence 24 hours before imaging. Cells were rinsed with buffer, and then maintained in buffer through imaging at 37° C using a spinning disk confocal microscope (Nikon Ti2 inverted microscope with a Yokogawa CSU-W1 confocal scanner). EM-CCD detectors (Andor, iXon 888 Ultra) and suitable dichroic and bandpass filters (from Semrock) were used for multicolor fluorescent imaging. Sequences of images shifting the sample in the axial direction (termed Z-stacks) were taken by manually identifying the bottom and top of the cell then obtaining fluorescent images for each color channel at 50 nm intervals. An exposure time of 200 ms was used for all CD9/CD63 images. An exposure time of 400 ms was used for images with the ER marker BFP-KDEL and the membrane dye.

### Manders Colocalization Analysis

This image analysis was done in Python, using the package numpy. Images were thresholded based on a manually determined cutoff and normalized so the sum of the pixel intensities was 1, making the images a probability density function rather than a matrix of fluorescence (photo-detection) counts. We then used the Manders test to compare the probability distributions. The Manders test compares the colocalization between two images (I_A_, I_B_)^35,36^. Two colocalization coefficients are calculated using the Manders test: M1 determines the fraction of I_A_ that is colocalized with I_B_, and M2 determines the fraction of I_B_ that is colocalized with I_A_.

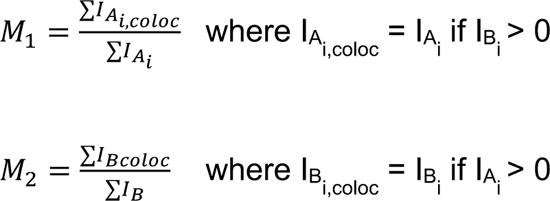

These two values allow quantifying the fraction of the total CD9 population colocalized with CD63, and the fraction of the total CD63 population colocalized with CD9. We expected these fractions to be different, making both of these quantities critical.

At the bottom of the cell (near the coverslip) and the top of the cell, the CD9 signal appeared across the entire area of the cell, which is consistent with an expression pattern we would expect if CD9 was associated with the cell membrane. This CD9 expression pattern confounded our ability to see whether CD9 and CD63 were colocalized at the top and bottom of the cell. Confocal slices have a finite depth of approximately 800 nm, so any CD63 within 800 nm of the cell surface would appear colocalized, even if the CD63 was just in close proximity. Clearly this was an artifact we wanted to avoid in our analysis. Therefore, we identified the top and bottom of the cells by identifying z-planes in which CD9 covered the entire area of the cell and excluded these z-planes from our colocalization analysis.

### Membrane Line Scan Analysis and Kolmogorov-Smirnov Test

The membrane line scan analysis was performed in Python with the packages numpy and scipy. Each color channel was thresholded separately (based on a manually identified cutoff). The location of the membrane was identified, and the x and y coordinates of each point along the membrane was recorded as follows. First, the centroid of the cell was identified by hand. Drawing a line from the centroid of the cell to a point on the membrane established an azimuthal angle *θ* and distance *r*(*θ*) to the membrane. We drew a line segment from the cell centroid through the point on the membrane so as to have an approximately equal distance within and beyond the cell membrane. The x and y coordinates of the points along this line were calculated as

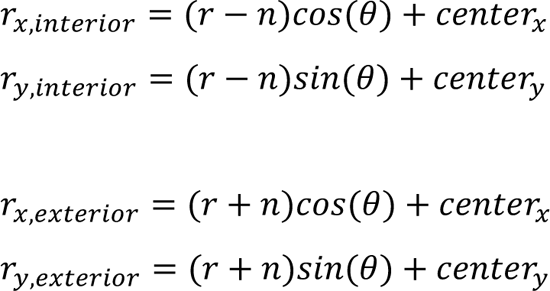

We repeated this for all points along the contour of the cell membrane, giving an array of values radially traversing the membrane at a specific membrane location. We then averaged the different radial arrays together and normalized so the distribution summed to 1. This allowed producing a single distribution for the entire contour of the membrane for each channel. Normalizing the arrays allowed comparing the distributions even when the intensities of the different channels varied. To study the relationship between the different labels (CD9, CD63, and the membrane), we applied the Kolmogorov-Smirnov test for each sample. The Kolmogorov-Smirnov test compares the difference between two probability distributions^37^. The smaller the KS coefficient, the closer or more similar the distributions are. Larger values of the KS coefficient (i.e., closer to 1) mean that the distributions are more different from one another.

## Acknowledgements

This work was supported by NIH awards R01 CA184494 (to MRR) and R35 GM136381 (to ARD) and a Big Ideas grant from the University of Chicago (to NFS). Elizabeth White acknowledges support from the National Institute of Biomedical Imaging and Bioengineering Grant (5T32EB009412). We further thank Dr. Kristin Luther (NanoView Biosciences, Boston, MA) for helping with the NanoView experiments and analysis.

