## Supplementary Information Text for "Volumetric microscopy of CD9 and CD63 reveals distinct subpopulations and novel structures of extracellular vesicles *in situ* in triple negative breast cancer cells"

Elizabeth D. White[1], Nykia D. Walker[2], Hannah Yi[3], Aaron R. Dinner[3,4*], Norbert F. Scherer[3,4*], Marsha Rich Rosner[2*]

[1] Graduate Program in Biophysics, University of Chicago, Chicago IL 60637

[2] Ben May Department for Cancer Research, University of Chicago, Chicago, IL 60637

[3] Department of Chemistry, University of Chicago, Chicago IL 60637

[4] Institute for Biophysical Dynamics, University of Chicago, Chicago IL 60637

*Corresponding Authors

Aaron Dinner,

Norbert Scherer,

Marsha Rosner,

### *In vitro* Studies of Extracellular Vesicles Secreted from the Cell Show a High Degree of CD9 and CD63 Co-localization

We used NanoView technology and quantified marker co-localization to verify whether or not EVs secreted from triple-negative breast cancer cells (BM1) have a large amount of co-localization between CD9 and CD63. NanoView captures EVs with a specific biomarker, in this case CD9, CD63, and CD81, on antibody coated chips, immunostains the EVs on each chip, and uses interferometric imaging to determine which other biomarkers (CD9, CD63, and CD81) are present on individual EVs[1–3] (Supplementary Figure 1). This technology is able to detect individual EVs, and is specifically calibrated to detect CD9, CD63, and CD81[2]. On the chip that captured CD9, 69% of the EVs also contained CD63. Similarly, on the chip that captured CD63, ~37% of the EVs also contained CD9 (Supplementary Figure 1).

### Observing Budding Extracellular Vesicles

There are multiple mechanisms through which EVs can be exported from the cell. We were able to observe the export of single CD9 and CD63 vesicles by identifying vesicles outside of, but close to, the cell membrane. However, a second mechanism of export is when vesicles bud directly from the cell membrane[4,5]. Both microvesicles and apoptotic bodies bud directly from the cell membrane; these two populations are differentiated primarily by their sizes[4]. Microvesicles are generally considered to be 100 nm -1 μm in diameter, while apoptotic bodies can be anywhere from 50 nm to 5 μm [4–6]. While the range of sizes are different, they do overlap.

CD63 is reported to be primarily in exosomes, which do not bud from the membrane[4,7]. Consistent with other reports, we did not find any examples of CD63 budding from the membrane. In contrast, CD9 is reported to be in microvesicles[4,5]. We found several examples of CD9 budding from the membrane, shown by vesicles forming from the membrane but not quite attached (see Supplementary Figure 2). Since the sizes of microvesicles and apoptotic bodies are overlapping, we were unable to determine which we were observing. Nevertheless, we can confidently say that we observed CD9 budding from the cell membrane.

### Verifying the Spatial Distributions of CD9 and CD63

One concern we had while conducting our study was whether the spatial patterns of CD9 and CD63 we observed were due to the inherent overexpression of the proteins from transfection. We controlled for this concern in two ways. First, we varied the amount of DNA used to transfect the cells over an order of magnitude (0.2 μg – 2 μg). When doing this, we saw no difference in the spatial distribution of the proteins (Supplementary Figure 3).

We immunostained CD9 and CD63 in BM1 cells to visualize the spatial distribution of the proteins at endogenous expression levels. The visual spatial distribution of the proteins matched what we observed in the transfected cells; CD9 was concentrated at the cell membrane, while CD63 was found primarily in the cytoplasm (Supplementary Figure 4A).

In addition to visualizing the spatial distribution of immunostained cells, we quantified the co-localization of the two proteins by calculating the Manders coefficient, the same way we did in transfected cells. There was even less co-localization between CD9 and CD63 in the immunostained cells (Supplementary Figure 4B). This confirmed that the separate spatial distribution of CD9 and CD63 was not due to overexpression of the proteins.

The last key observation we wanted to verify in cells with endogenous expression levels of CD9 and CD63 was to see whether the shell-like structures in which CD63 encapsulates CD9 we saw in transfected cells were present. We did observe these donut-like structures (Supplementary Figure 5).

Varying the transfection amounts of CD9 and CD63 and staining CD9 and CD63 to observe their endogenous expression levels, were essential controls to verify that the spatial distribution we observed was not due to over expression of the proteins. Replicating the analysis for the co-localization of CD9 and CD63 in the stained cells further reinforced that the lack of co-localization between the two proteins is accurate. Observing the same donut-like structure of CD9 and CD63 in the endogenously labeled cells also confirmed that we were not altering the spatial distribution of CD9 and CD63 when transfecting the cells.

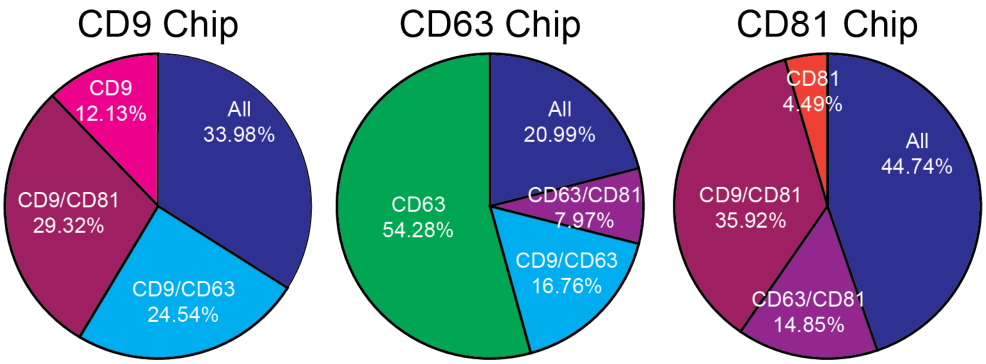


**Fig. S1.** *In vitro* analysis of extracellular vesicles shows significant co-localization of CD9, CD63, and CD81. EVs captured by CD9 (left, N = 17,120), CD63 (middle, N = 5,694), and CD81 (right, N = 4,259) are labeled with other antibodies. Percentage of EVs containing different combinations of antibodies are calculated and shown in white text.

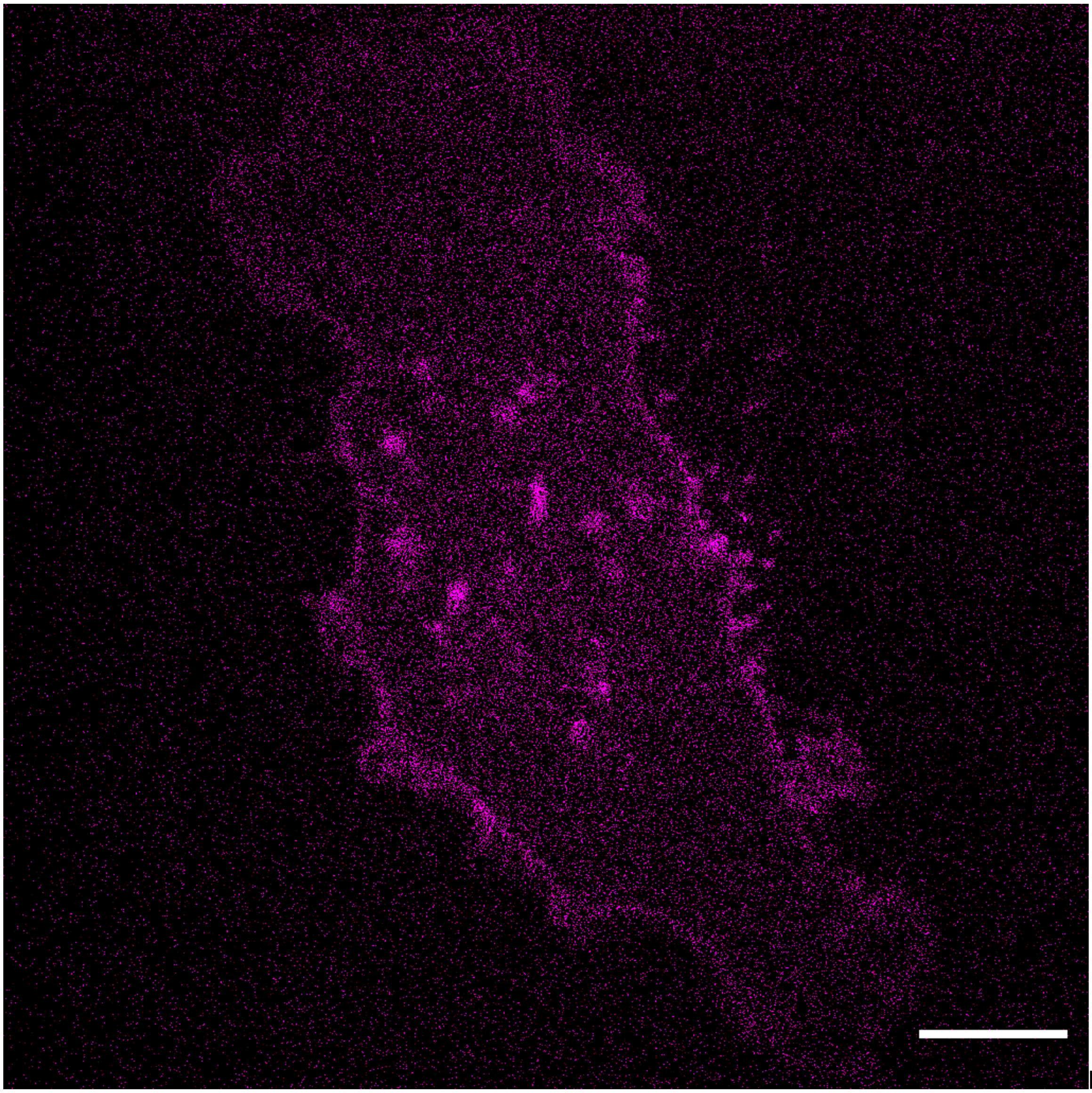

**Fig. S2.** Evidence of CD9 budding. Cells were transfected with 0.5 µg CD9-mCherry DNA. White boxes indicate regions of CD9 budding. Scale bar: 10 µm.

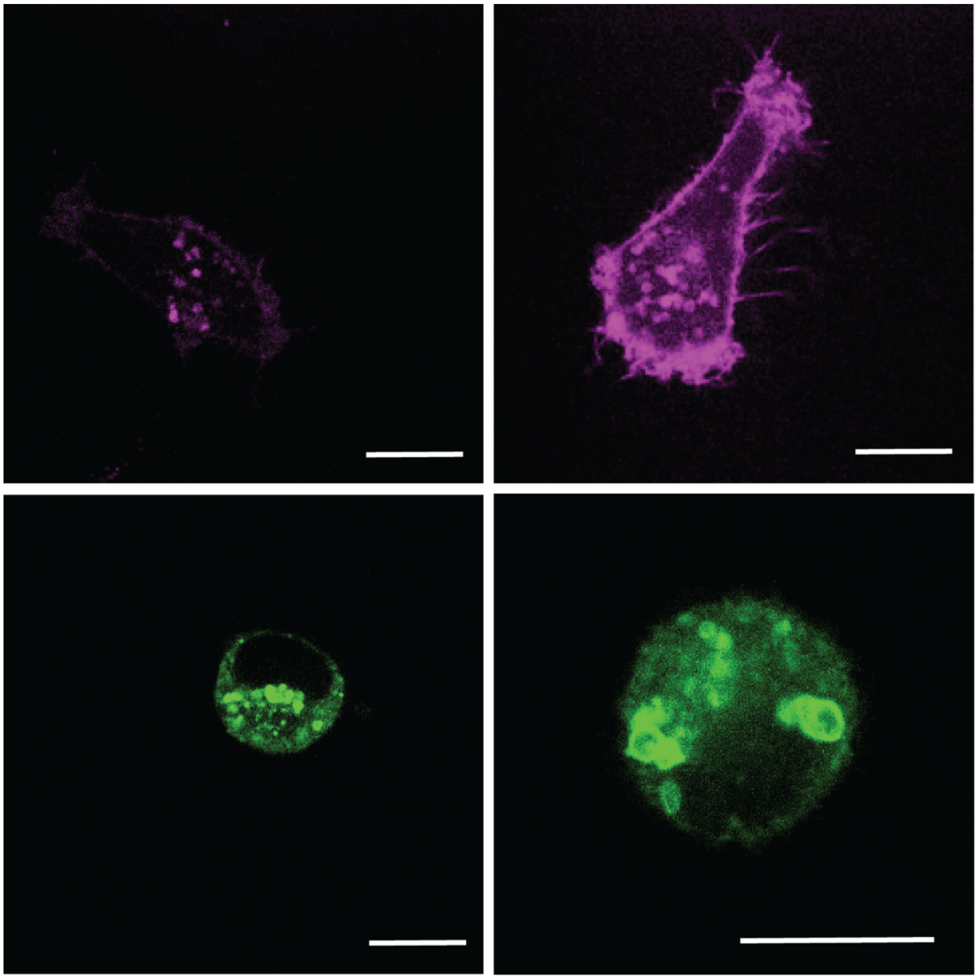

**Fig. S3.** Cells transfected with different amounts of CD9-mCherry and CD63-GFP have similar spatial distributions. Top: BM1 cells transfected with 0.2 μg (left) and 2 μg (right) CD9-mCherry. Bottom: BM1 cells transfected with 0.2 μg (left) and 2 μg (right CD63-GFP.

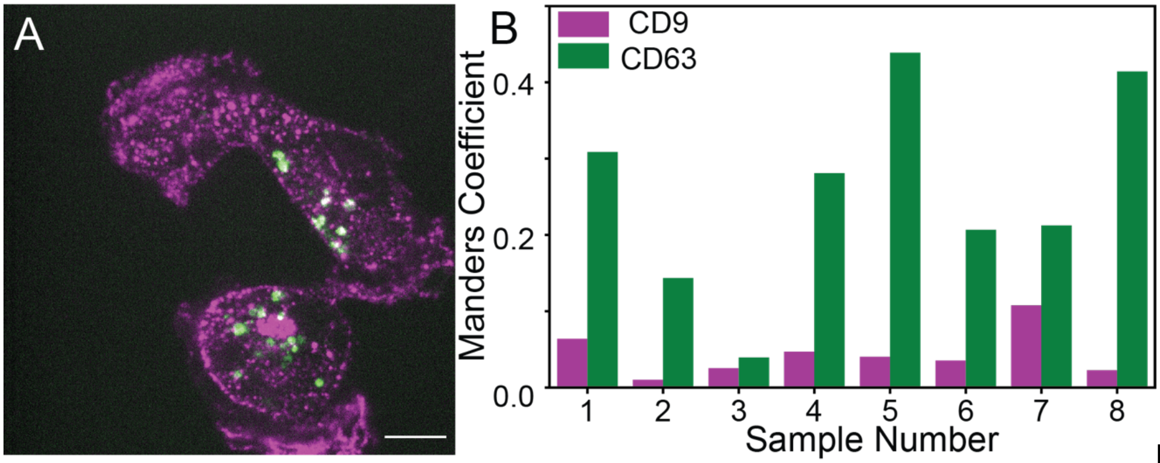


**Fig. S4.** CD9 and CD63 are spatially distinct in immunostained cells. A: BM1 cells stained with CD9 (magenta) and CD63 (green) show that they two labels rarely overlap. B: Manders' coefficient for 8 samples was calculated, showing little overlap between CD9 and CD63.

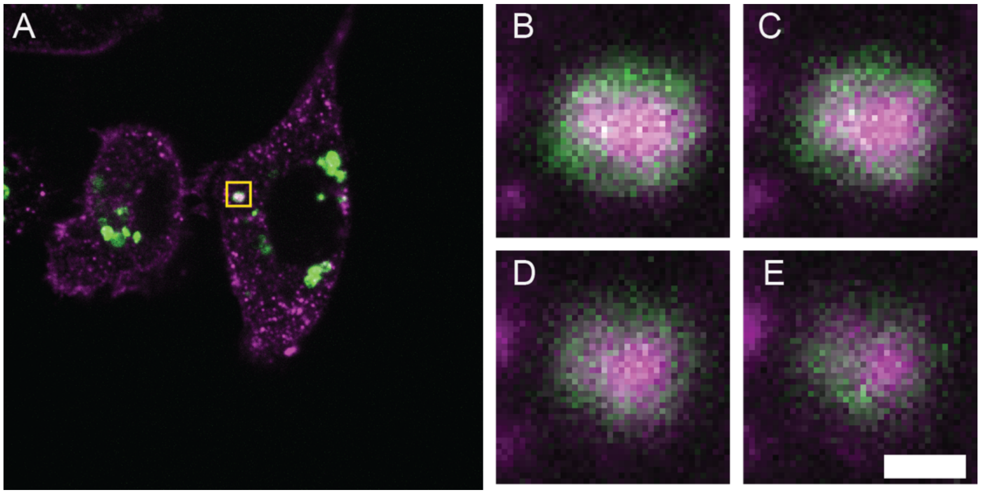


**Fig. S5.** Immunostaining CD9 (magenta) and CD63 (green) reveals similar spatial distributions as transfecting CD9-mCherry and CD63-GFP. White box highlights a donut-like structures that are also observed in other cells. Scale bar: 10 μm.

**Movie S1 (separate file).** Z-stack of CD9-mCherry in BM1 cells. Scale bar: 10 µm. Z-step: 50 nm.

**Movie S2 (separate file).** Z-stack of CD63-GFP in BM1 cells. Scale bar: 10 µm. Z-step: 50 nm.

**Movie S3 (separate file).** Z-stack of CD9-mCherry and CD63-GFP in BM1 cells. Scale bar: 10 µm. Z-step: 50 nm.
